# Electron tomography visualization of HIV-1 fusion with target cells using fusion inhibitors to trap the pre-hairpin intermediate

**DOI:** 10.1101/2020.04.29.068775

**Authors:** Mark S. Ladinsky, Priyanthi N.P. Gnanapragasam, Zhi Yang, Anthony P. West, Michael S Kay, Pamela J. Bjorkman

## Abstract

Fusion of HIV-1 with the membrane of its target cell, an obligate first step in virus infectivity, is mediated by binding of the viral envelope (Env) spike protein to its receptors, CD4 and CCR5/CXCR4, on the cell surface. The process of viral fusion appears to be fast compared with viral egress and has not been visualized by electron microscopy (EM). To capture fusion events for EM, the process must be slowed or stopped by trapping Env-receptor binding at an intermediate stage. Here we describe using fusion inhibitors to trap HIV-1 virions attached to target cells by Envs in an extended pre-hairpin intermediate state. Electron tomography revealed HIV-1 virions bound to TZM-bl cells by 2-4 narrow spokes, with slightly more spokes present when evaluated with mutant virions that lacked the Env cytoplasmic tail. These results represent the first direct visualization of the hypothesized pre-hairpin intermediate and improve our understanding of Env-mediated HIV-1 fusion and infection of host cells.

## Introduction

The first step of HIV-1 entry into a host target cell, fusion between the viral and target cell membranes, is mediated by the viral envelope spike protein (Env). HIV-1 Env is a trimeric glycoprotein comprising three gp120 subunits that contain host receptor binding sites and three gp41 subunits that include the fusion peptide and membrane-spanning regions. Binding of the primary receptor CD4 to gp120 triggers conformational changes that expose a binding site for co-receptor (CCR5 or CXCR4). Coreceptor binding results in further conformational changes within gp41 that promote release of the hydrophobic fusion peptide, its insertion into the host cell membrane, and subsequent fusion of the host cell and viral membrane bilayers [1].

Structural studies relevant to understanding Env-mediated membrane fusion include X-ray and single-particle cryo-EM structures of soluble native-like Env trimers in the closed (pre-fusion) conformation [2], CD4-bound open trimers in which the co-receptor binding site on the third hypervariable loop (V3) of gp120 is exposed by V1V2 loop rearrangement [3–6], a gp120 monomeric core-CD4-CCR5 complex [7], and a post-fusion gp41 six-helical bundle formed by an a-helical trimeric coiled coil from the gp41 N-trimer region surrounded by three helices from the C-peptide region [8, 9] (Figure 1a). Prior to membrane fusion and formation of the postfusion gp41 helical bundle, the viral and host cell membranes are hypothesized to be linked by an extended pre-hairpin intermediate in which insertion of the gp41 fusion peptide into the host cell membrane exposes the N-trimer (HR1) region of gp41 [10]. Formation of the six-helical bundle and subsequent fusion can be inhibited by targeting the N-trimer region with C-peptide-based inhibitors; e.g., the fusion inhibitor T20 (enfuvirtide [Fuzeon]) [11], T1249, a more potent derivative of T20 [12], and a highly potent trimeric D-peptide (CPT31) [13], or with anti-gp41 antibodies such as D5 [14] (Figure 1a).

**Figure 1.**
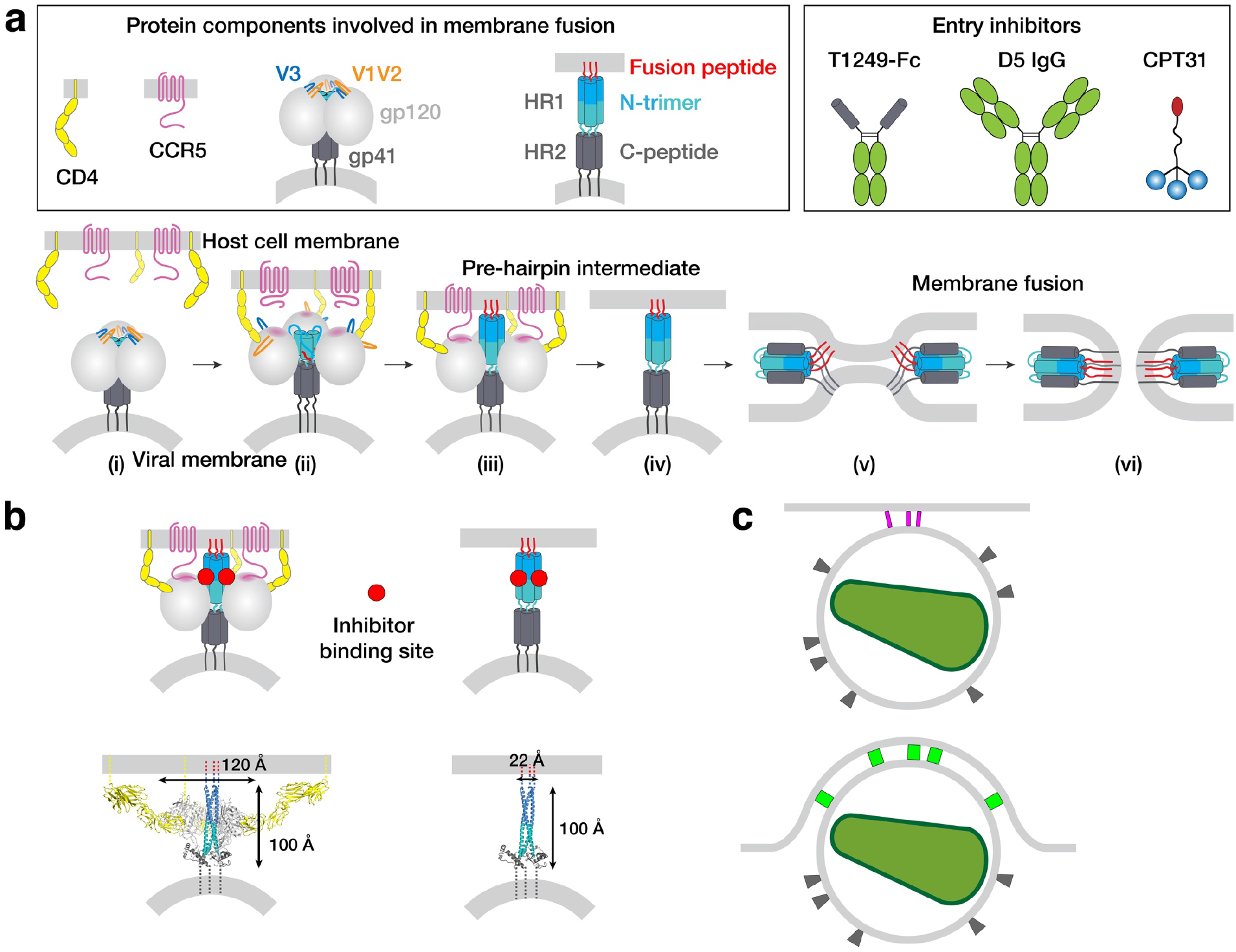
HIV-1 Env-mediated fusion between viral and host cell membranes. **a**, Top: Schematics of host receptors, HIV-1 Env trimer, pre-hairpin intermediate, and fusion inhibitors. Bottom: steps in fusion: (i) Closed, prefusion structure of HIV-1 Env trimer in which the V1V2 loops (orange) occlude the coreceptor binding site on V3 (blue) (e.g., PDB code 5CEZ). The Env trimer is embedded in the viral membrane, while the host receptor (CD4) and coreceptor (CCR5) are embedded in the target cell membrane. (ii) CD4-bound open HIV-1 Env trimer in which V1V2 loops have been displaced to expose the coreceptor binding site on V3 (e.g., PDB 6U0L). (iii) Hypothetical CD4- and CCR5-bound open Env trimer with rearrangements of gp41 N-trimer/HR1 to form a pre-hairpin intermediate structure that is linked to the target cell membrane by the gp41 fusion peptide (red). (iv) Hypothetical pre-hairpin intermediate formed by gp41 trimer after shedding of gp120s. (v-vi) Formation of the post-fusion gp41 six-helical bundle (e.g., PDB 1GZL) that juxtaposes the host cell and viral membranes (step v) for subsequent membrane fusion (step vi). **b**, Approximate binding sites (red circles) for fusion inhibitors shown on schematics of steps iii and iv (panel a). Entry inhibitor binding sites might be partially sterically occluded for binding to the T1249-Fc or D5 fusion inhibitors. Schematics shown above as models are from PDB codes 6U0L and 1AIK with approximate dimensions indicated. **c**, Schematic illustrating why fewer HIV-1 Envs might be involved in attaching to a target cell when the attachment site is flat versus a concave surface. Top: attachment site (described here) formed during a 37°C incubation of virions, target cells, and a fusion inhibitor. Bottom: attachment site (described in [25]) formed in the absence of a fusion inhibitor when virions and target cells were incubated in a temperature jump protocol (4°C incubation followed by warming to 37°C).

Visualizing the pre-hairpin intermediate that joins the host and viral membranes has not been straightforward. Despite 3-D imaging by electron tomography (ET) of HIV-1 infection of cultured cells [15–18] and tissues [19–22], viruses caught in the act of fusion have not been unambiguously found. In our ET imaging of HIV-1–infected humanized mouse tissues, we have identified hundreds of budding virions at various stages of egress and thousands of free mature and immature virions [19–22], but not a single example of a virus attached to a host cell via a pre-hairpin intermediate or in the process of fusing its membrane with the target cell membrane. The absence of observed viral fusion events might be explained if fusion is a fast process compared with viral budding; thus when cells or tissues are immobilized for EM or ET, the relatively slow process of viral budding would be more easily captured compared with the faster process of fusion. We assume that fusion could theoretically be observed if a virus were caught at exactly the right time, but this might require examining thousands or millions of images.

Here we report visualizing the pre-hairpin intermediate by ET after treatment of HIV-1–exposed target cells with inhibitors of six-helix bundle formation that bind the N-trimer region of gp41 that is exposed during the fusion process. Using optimally preserved samples for ET with a nominal resolution ~7 nm, we found >100 examples of HIV-1 virions linked to TZM-bl target cells by 2-4 narrow rods of density (spokes) in inhibitor-treated samples, but none in untreated or control-treated samples. The approximate dimensions of the majority of the spokes matched models of gp41-only pre-hairpin intermediates in which the Env gp120 subunit had been shed. The average number of observed spokes connecting a virion to a target cell increased slightly when using a virus containing an Env with a cytoplasmic tail deletion, suggesting that the increased lateral mobility of cytoplasmic tail-deleted Envs in the viral membrane [23, 24] allowed more Envs to join the interaction with the target cell. We discuss the implications of these studies for understanding HIV-1 Env-mediated membrane fusion and how these results differ from a previous ET study of the “entry claw” that is formed upon HIV-1 or SIV interactions with target cells [25].

## Results

### Experimental design

A previous study used ET to visualize HIV-1 and SIV virions in contact with target cells after promoting a temperature-arrested state [26] in which viruses can remain attached to cells prior to fusion [25]. For that study, target cells were incubated with virus at 4°C to allow binding but not fusion, warmed to 37°C, and then fixed after incubations ranging from 15 min to 3 h [25]. At all time points after warming, viruses were found attached to target cells by a cluster of 5-7 “rods,” each ~100 Å long and ~100 Å wide. The fact that the attachment structure was not found when the viruses and target cells were incubated in the presence of C34, a gp41 N-trimer–targeting C-peptide inhibitor related to T20 [25], suggests that the rod structure that was trapped during the temperature-arrested state did not involve the pre-hairpin intermediate.

We hypothesized that addition of an HIV-1 fusion inhibitor that binds to the exposed gp41 N-trimer after host cell receptor and coreceptor binding would slow or stop virus-host cell membrane fusion such that we could visualize pre-hairpin intermediate structures by ET (Figure 1b). In addition, incubating with a fusion inhibitor at 37°C obviated the need for a 4°C incubation of virus and target cells, which we reasoned was desirable since low temperatures alter membrane fluidity [27–29], which could affect one or more steps in membrane fusion. Since target cells for HIV-1 are several microns in height, much thicker than the 0.5-1 μm limit for cryo-ET [30], we used stained, plastic-embedded samples that could be cut into 300-400 nm sections using a microtome, and then examined the samples in 3-D using ET. Furthermore, because attached virions were rare, it was an advantage that radiation damage is minimal in plastic sections [31], thus more cells could be assayed in plastic sections than in samples prepared by cryo-ET methods (e.g., by examining thin leading edges of cells or using focused-ion-beam milling [32] to prepare a sufficiently thin sample), thus allowing for statistically-significant observations of virion attachment events. We prepared samples by light fixation followed by high-pressure freezing/freeze substitution fixation (HPF-FSF) instead of the traditional chemical fixation protocol used previously [25] because HPF vitrifies cells at ~10,000°/sec, stopping all cellular movement within msec and allowing optimal preservation of ultrastructural features [33–36]. By contrast, chemical fixation immobilizes elements in the cell at different rates, and movement and rearrangement of transmembrane proteins may continue even in the presence of aldehyde fixatives [37–39]. Following HPF-FSF, samples were plastic embedded and stained with uranyl acetate and lead citrate as described in our previous ET studies of HIV-1 in infected tissues [20–22]. Since biosafety requirements for the current study necessitated the use of HIV-1 pseudoviruses instead of infectious HIV-1, we verified that the ultrastructure of HIV-1 pseudoviruses, including approximate numbers and dimensions of Env trimer spikes and the presence of collapsed (in mature virions) versus C-shaped (in immature virions) cores [15, 40–42], was preserved during the fixation, embedding, and staining procedures (Supplementary Figure 1), consistent with our previous publications involving ET of infectious HIV-1 in tissue samples [20–22]. These results are also consistent with previous direct comparisons of tomograms of stained and plastic-embedded versus unstained and cryopreserved SIV virions [25].

We characterized three fusion inhibitors of different sizes and potencies that target the exposed gp41 N-trimer region of the pre-hairpin intermediate for attempts to visualize the pre-hairpin intermediate: T1249-Fc, a C-peptide–based inhibitor that we linked to human Fc (MW = 65 kDa), D5 IgG (MW = 150 kDa) [14], and CPT31, a high-affinity D-peptide inhibitor linked to cholesterol (MW = 9 kDa) [13, 43] (Figure 1a; Supplementary Figure 2a). We measured their neutralization potencies using in vitro HIV-1 pseudovirus neutralization assays [44] against the SC4226618 and 6535 viral strinas. We found potencies ranging from 50% inhibitory concentration (IC_50_) values of ~0.13 ng/mL for CPT31 to ≥40 μg/mL for D5 IgG (Supplementary Figure 2a). T1249-Fc exhibited intermediate potencies (IC_50s_ = 0.99 μg/mL; 17 μg/mL) (Supplementary Figure 2a), higher than IC_50s_ measured for T1249 peptide alone, consistent with limited steric accessibility resulting in decreased potencies for larger fusion inhibitors [45].

We conducted ET experiments by first incubating TZM-bl cells, a HeLa cell line that stably expresses high levels of human CD4 and coreceptors CCR5 and CXCR4 [46], with 130 μg/mL of inhibitor (either T1249-Fc, D5, or CPT31) and ~5000 TCID_50_/mL of HIV-1 pseudovirus at 37°C for 2, 4, or 48 hours, followed by HPF, FSF, plastic embedding, sectioning, and visualization by ET. In order to verify that results were not dependent upon a particular viral strain, we conducted experiments using pseudoviruses derived from two primary isolate HIV-1 strains: SC4226618 (Tier 2) and 6535 (Tier 1B) [47], chosen for their sensitivity to the fusion inhibitors (Supplementary Figure 2a) and because we had both wild-type and Env cytoplasmic tail-deleted forms of the 6535 pseudovirus. To identify attached virions by EM (Figure 2, Supplementary Figure 3), the peripheries of TZM-bl cells were scanned to locate roughly spherical objects with diameters ~100 nm that were near a cell surface. Regions of interest were then examined at higher magnification and at tilts of 0°, 35° and −35° to verify that the objects were spherical, as expected for an HIV-1 virion. Potential virions were then observed through a defocus series to detect core structures found inside authentic virions: i.e., a bullet-shaped core in mature HIV-1 and a C-shaped core in immature HIV-1 [40, 42]. Once verified as a virion, tilt series for 3-D reconstructions were collected. Control experiments in which pseudovirus and TZM-bl cells were incubated without inhibitor, with an irrelevant IgG, or with a low concentration of inhibitor, were prepared and analyzed in the same way.

**Figure 2.**
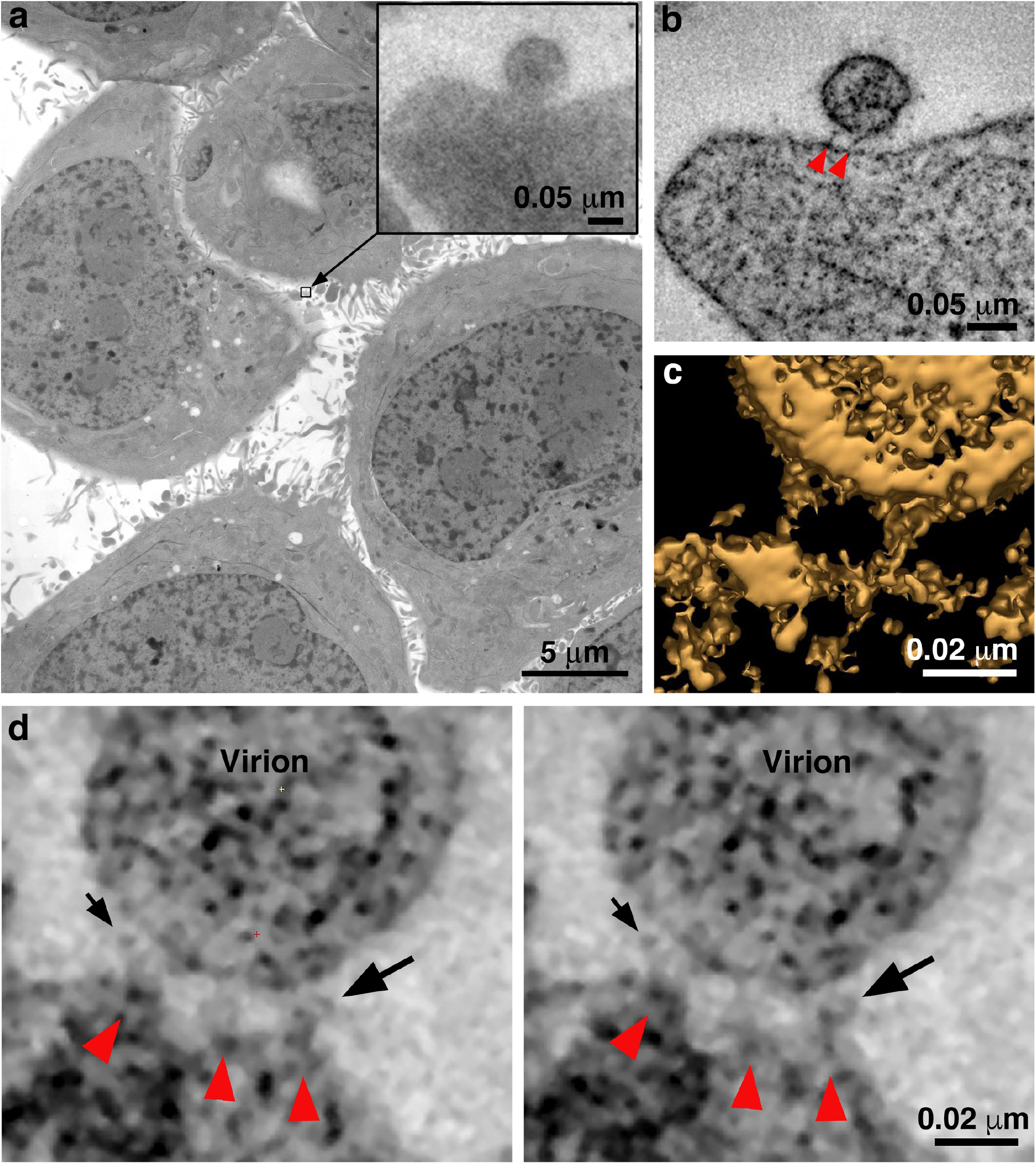
Identification of attached HIV-1 virions. **a**, Montaged projection overview of a field of cultured TZM-bl cells from a 400 nm section. Note extensive blebbing and surface projections that are typical of the cell type. Inset: Projection detail of a HIV-1 virion adjacent to TZM-bl cell surface. **b,** Slice (5.6 nm) from a tomographic reconstruction of the virion shown in the inset of panel a (from a dataset collected with the T1249-Fc inhibitor). The bullet-shaped core identifies the particle as mature HIV-1 (see also Supplementary Figure 3). Two pre-hairpin intermediate “spokes” (red arrowheads) attach the virion to the cell surface. **c**, 3-D isosurface rendering of the spokes shown in panel b. **d**, Examples of extra densities observed in some data sets collected using the D5 IgG inhibitor. These appear as “hook-like” structures projecting from the sides of spokes, adjacent to the virion surface, which are visible in two sequential tomographic slices (small and large black arrows). Extra densities may represent portions of D5 IgGs attached to the prehairpin intermediate. Similar densities were not seen in experiments with the T1249-Fc or CPT31 inhibitors.

TZM-bl cells are contaminated with ecotropic murine leukemia virus [48], which does not affect their use for HIV-1 in vitro neutralization assays [49]. In our surveys of TZM-bl cells incubated in the presence or absence of fusion inhibitors, we occasionally observed budding MLV virions (Supplementary Figure 4). As MLV serves as a control for non-specific inhibition in TZM-bl–based HIV-1 in vitro neutralization assays [50], the fusion inhibitors used in our experiments are known to have no effect on MLV fusion, thus contaminating MLV virions were not captured during fusion in our experiments.

### Entry inhibitor-treated virions are attached to target cells by 2-3 narrow spokes

In surveys of TZM-bl cells treated with either SC4226618 or 6535 pseudovirus and 130 μg/mL of any of the three fusion inhibitors, we found virions that were attached to the surface of a TZM-bl cell by several (usually 2 or 3) narrow densities. To verify that the attached virions resulted from treatment with a fusion inhibitor, we analyzed control experiments in which we incubated TZM-bl cells and pseudovirus with either no inhibitor, with an irrelevant Fc-containing protein (Z004, an anti-Zika virus IgG [51]), or with the T1249-Fc inhibitor at a concentration equivalent to 0.01x of its neutralization potency (i.e., its IC_50_ value) (Supplementary Figure 2a). In examinations of >100 cells, we found no attached virions and also very few virions that were within a distance that could accommodate attachment. We used ET to examine the few cases (<20) in which we found virions adjacent to cells, which confirmed that none of the virions were attached to a host cell membrane (Supplementary Figure 5).

The approximate dimensions of the densities found for virions attached to TZM-bl cells in the presence of a fusion inhibitor were 11-15 nm in length and ~2-4 nm in width (Figure 3), thus we refer to the densities as “spokes” to distinguish them from the wider densities described as “rods” in the previous ET study of virions attached to cells [25]. In some samples incubated with the largest inhibitor (D5 IgG), we occasionally found densities adjacent to one or more spokes that could represent the bound IgG (Figure 2d). We found virions attached with spokes to 10-30% of TZM-bl cells that were examined, usually located on the thin leading edges of cells. Most cells showed only one attached virion per 400 nm section, but occasional cells exhibited 3-5 attached virions. The attachment sites were generally flat, as opposed to the target cell exhibiting a concave surface corresponding to the circumference of the virion as previously described [25], with distances of ~7 nm to ~15 nm between spokes. The majority of attached virions were mature, as identified by their bullet-shaped cores (Figures 2, 3), but a minor subset of attached virions (~5%) were immature. Figure 2 shows overview images and 2-D tomographic slices from 3-D reconstructions of fusion inhibitor-treated virions attached to target cells; see also a gallery of examples in Supplementary Figure 6. Attachment densities were sometimes located in different planes so they are not always visible in any given 2-D tomographic slice, thus we identified them in 3-D as shown in Supplementary Movie 1.

**Figure 3.**
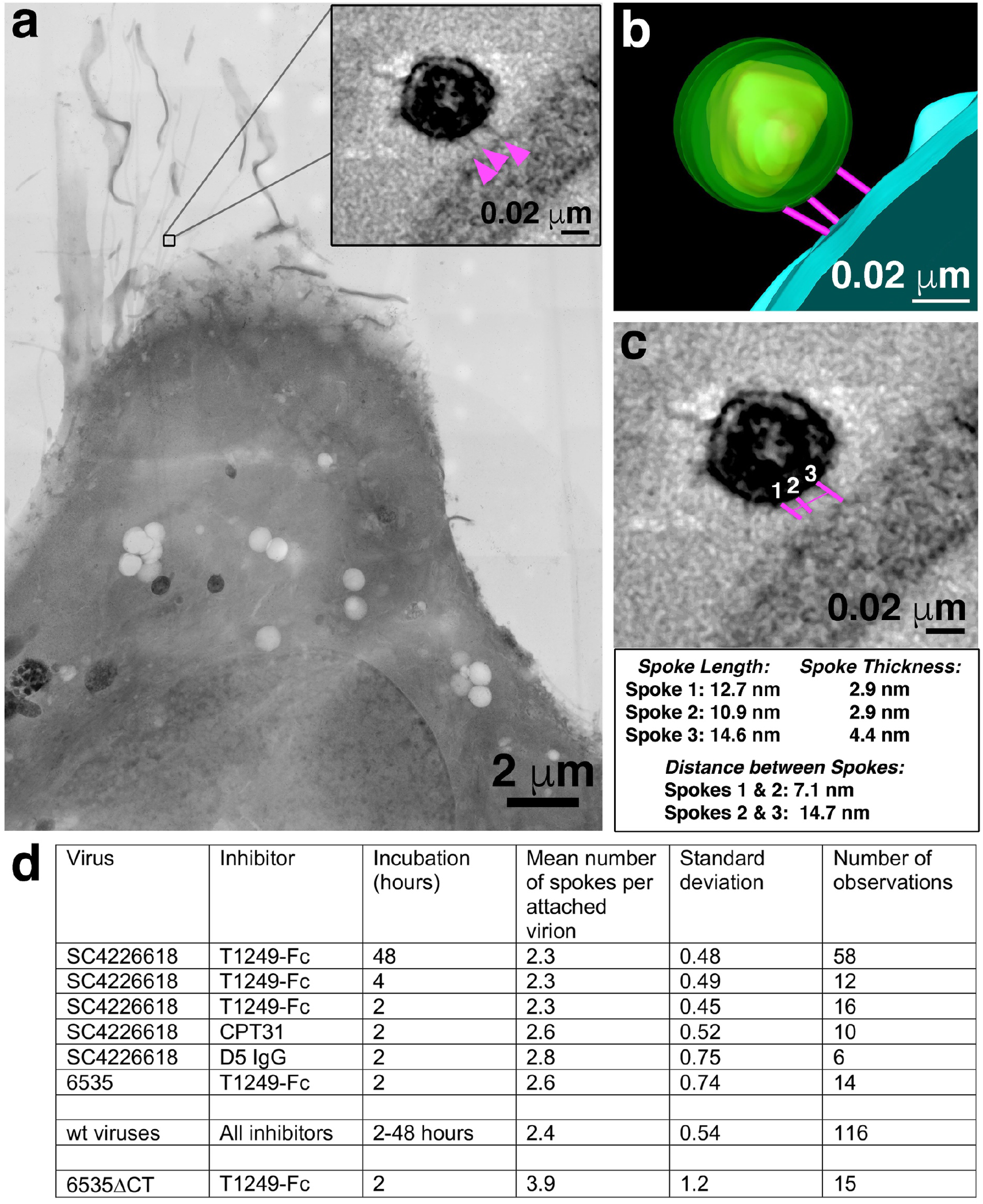
Characteristics of virions attached to target cells in the presence of a fusion inhibitor. **a**, 2-D projection image of TZM-bl cell incubated with SC4226618 pseudovirions in the presence of the T1249-Fc fusion inhibitor for 2 hours at 37°C. Inset shows a tomographic slice of the attached virion indicated by a box with attachment spokes indicated by magenta arrowheads. **b**, 3-D model from tomogram of attachment site shown in panel a inset. **c**, Tomographic slice of attached virion from panel b with approximate measurements of spoke length, width, and inter-spoke distances. **d**, Summary of mean, standard deviation, and number of observations for spokes at attachment sites under different experimental conditions. See also gallery of images in Supplementary Figure 6.

Despite using inhibitors of different sizes and at concentrations varying from 3.2- to 10^6^-fold above their IC_50_ values for neutralization of the SC4226618 pseudovirus (Supplementary Figure 2a), we found no systematic differences in the numbers of attached virions per cell, their locations on TZM-bl cells, or the numbers and dimensions of spokes per attached virion. In addition, no systematic differences were observed as a function of which fusion inhibitor or pseudovirus was included in the incubation or the length of the 37°C incubation (Figure 3d; Extended Data Table 1). Although a 48-hour incubation at 37°C should result in a substantial loss of virus infectivity, the T1249-Fc incubation for 48 hours condition yielded an equivalent number of attached virions (Figure 3d) and similar spoke structures as the 2- and 4-hour incubations (Supplementary Figure 6). This finding is rationalized by calculations and infectivity measurements showing that the 48-hour incubation conditions contained several million infectious virions (Supplementary Figure 2b), enough to account for the observed attached virions.

### A cytoplasmic tail deletion virus forms attachment structures with more spokes

The finding of only 2-3 spokes per attached virion implies that not all of the ~14 Envs per HIV-1 virion [52–56] are involved in the fusion process. Indeed, Env trimers that did not participate in attachment were sometimes observed in tomographic slices (Figure 4). We hypothesized that more Envs might join in the attachment site if we used a virus in which Envs could exhibit increased lateral mobility in the viral membrane to allow Env diffusion to the attachment site. Increased Env mobility in the viral membrane has been described in viruses with an Env cytoplasmic tail deletion that eliminates interactions with the viral matrix protein [23, 24]. To evaluate potential effects of increased lateral mobility, we used a pseudovirus containing an Env with a cytoplasmic tail deletion (6535-ΔCT) and compared its attachment sites with target cells when incubated with the T1249-Fc fusion inhibitor to those of wild-type 6535 and SC4226618 pseudoviruses when incubated under the same conditions (Figure 5).

**Figure 4.**
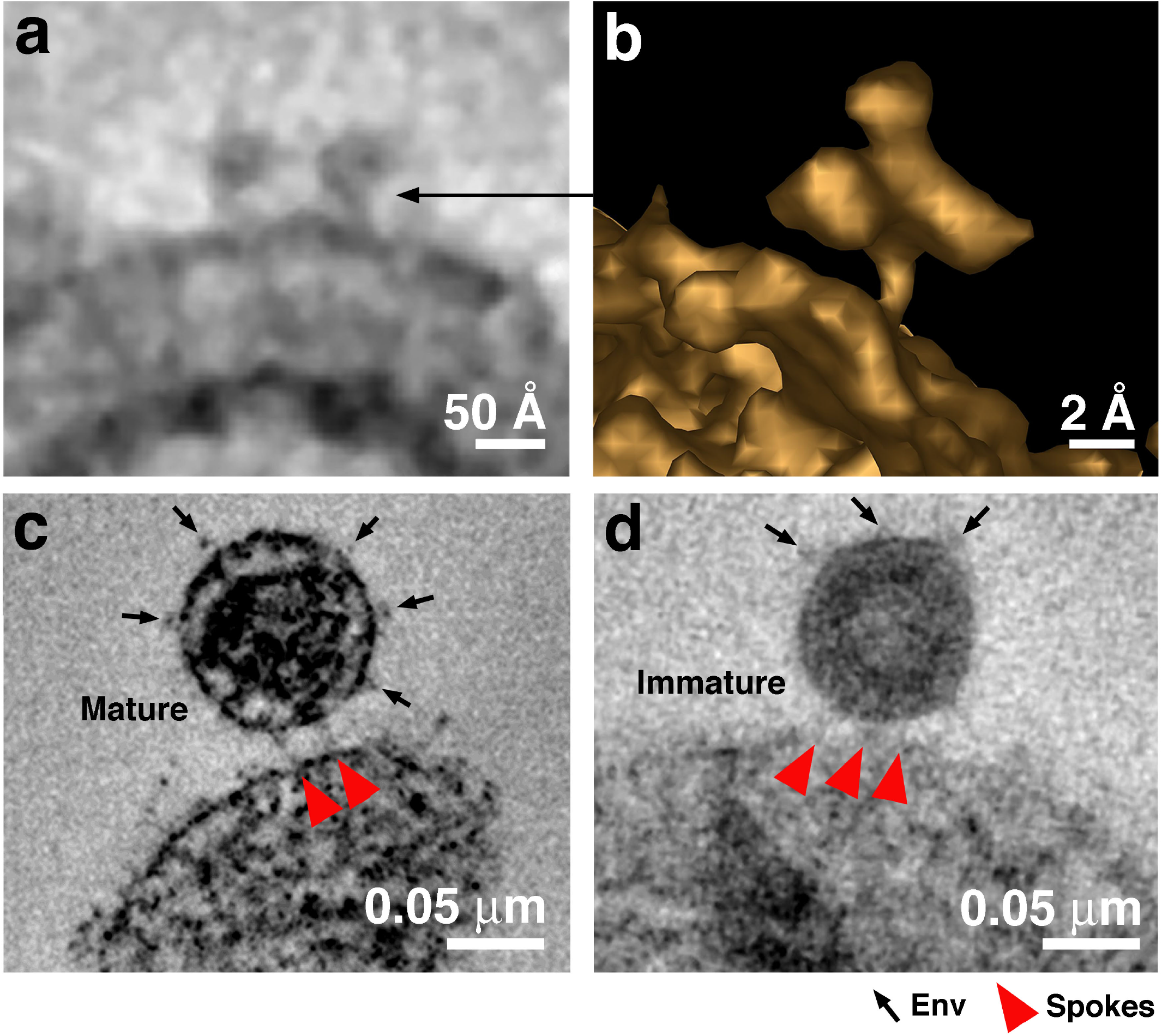
Free Env trimers can be visualized on attached virions. **a**, Example of densities observed for free HIV-1 Env trimers in a tomographic slice. **b**, 3-D isosurface rendering of an individual free Env trimer. **c-d**, Examples of attached mature (panel c) and immature (panel d) virions with free Env spikes distant from the attachment site indicated by arrows. Note that free Env trimer densities and spokes at an attachment site were only rarely optimally visualized in a single tomographic slice, thus Env trimers and spokes were identified from 3-D tomograms rather than 2-D tomographic slices.

**Figure 5.**
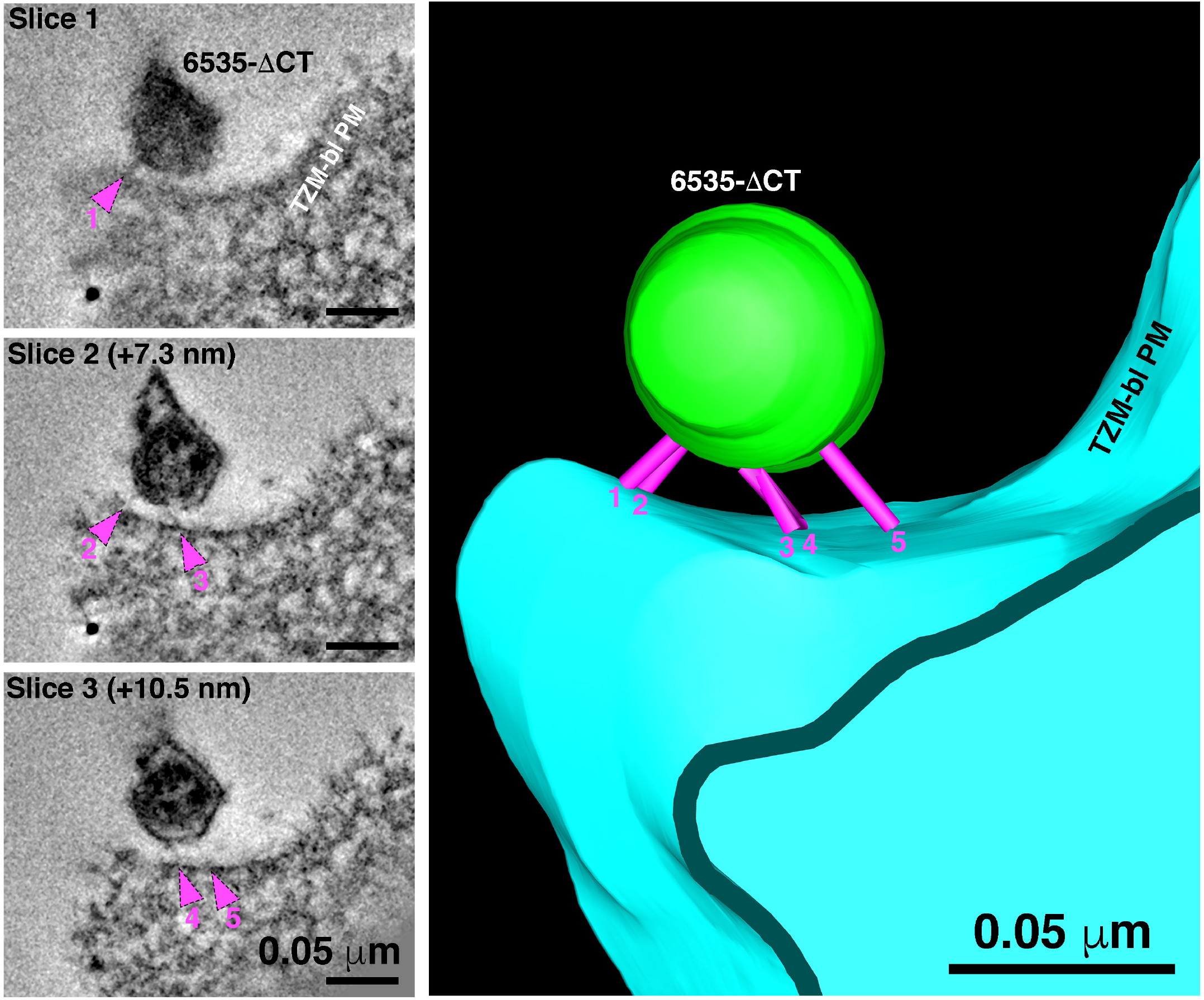
6535ΔCT pseudoviruses are often attached to target cells with more than 2-3 spokes. An example shows a 6535ΔCT virion attached to the plasma membrane of a TZM-bl cell (cell and virus were treated with the T1249-Fc fusion inhibitor for 2 hours). Viewing the tomogram in a series of slices through the reconstruction (left panels) reveals 5 distinct spokes (magenta arrowheads) at different positions on the virus’ surface. 3-D model of the reconstruction (right) displays the 5 spokes (magenta) attaching the virion (green) to the TZM-bl plasma membrane (PM) (blue).

We first confirmed that attachment sites formed by wild-type 6535 and SC4226618 pseudoviruses were indistinguishable in terms of their locations and numbers of observed spokes (Figure 3). We then gathered data for 6535-ΔCT attachment sites, finding a small, but statistically significant (*p* = 3 x 10^-4^) increase in the mean number of spokes: 3.9 +/- 1.2 (*n* = 15) spokes for 6535-ΔCT as compared to 2.4 +/- 0.6 (*n* = 116) spokes for wild-type pseudoviruses (Figures 3d, 5).

## Discussion

Despite a wealth of information about the pre- and post-fusion structures of HIV-1 Env and Env’s interactions with host receptors [2–9, 54], the architecture of the virus-host cell contact and the structure of Env during the act of fusion have remained elusive. Because HIV-1 virions have not been visualized during the act of fusion, we assumed that viral fusion is a fast process that has yet to be captured by electron microscopy imaging of either cultured cells or tissues. In order to visualize the hypothesized pre-hairpin intermediate structure in which the gp41 transmembrane region and the fusion peptide link the viral and target cell membranes [10], we incubated HIV-1 fusion inhibitors that bind to the N-trimer region of gp41 that is exposed upon Env binding to the host receptor and coreceptor on target cells. We chose to use diverse fusion inhibitors to trap the hypothesized pre-hairpin intermediate structure linking the viral and host cell membranes (Figure 1a) and to avoid a temperature jump protocol [25] that could result in unanticipated effects on membranes; for example, membrane trafficking events can be altered or stopped at decreased, non-physiological temperatures, and structures associated with them, such as Golgi cisternae, are perturbed and noticeably altered relative to physiological conditions [57].

We chose three fusion inhibitors with strong, medium, and weak neutralization potencies (CPT31, T1249-Fc, D5 IgG, respectively), incubated them at 37°C with virus and target cells for variable times, and examined them in 3-D by ET. For all experimental conditions, we found virions linked to the surface of target cells by several (mean = 2.4 +/- 0.5; *n* = 116) narrow spokes (Figure 3d), thus all conditions contained sufficient numbers of virions and inhibitors to form stable attachments to target cells. By contrast, in control experiments with either no inhibitor, an irrelevant protein, or inhibitor added at a concentration equal to 1% of its neutralization IC_50_ value, we did not observe virions attached to target cells. Based on current understanding of how fusion inhibitors prevent fusion [10, 58] (Figure 1a,b), we conclude that the attachment densities represent Env trimers in the pre-hairpin intermediate conformation that link viral and target cell membranes (Figure 1a).

Having trapped virions in the act of fusion, we can use information gathered from tomograms to address mechanistic and structural details of fusion between viral and target cell membranes. First, the results suggest that only 2 to 3 (and occasionally 4) Envs participate in the reaction with target cells that is inhibited by a fusion inhibitor. This number represents a minority of the Env trimers on HIV-1, even considering that HIV-1 includes a unusually low number of spikes per virion: an average of ~14 with a range from 4 – 35 [52–56] (By comparison, similarly-sized influenza A virions contain ~450 spikes [59]). STED microscopy studies suggested that HIV-1 spikes rearrange from a random distribution on immature virions to a cluster on mature virions [60, 61]. However, interspike distance distributions on mature virions derived from independent cryo-ET reconstructions revealed random spike distributions rather than a single cluster of spikes [53, 54]. Our finding of only 2-3 spokes per virion-target cell attachment site (Figure 3d Extended Data Table 1) is also consistent with random spike distributions on mature virions since more attachment spokes would be expected if Envs are clustered. In addition, although free Env trimers on HIV-1 virions are difficult to identify conclusively by ET using stained, plastic-embedded samples, we sometimes found Env spikes on the opposite surface of attached mature virions as the attachment structures (Figure 4c,d), consistent with the assumption that only two or three Envs are usually close enough to each other to participate in binding to host cell receptors and that slow lateral diffusion of HIV-1 Envs within the virion bilayer prohibits recruitment of additional Envs into the attachment structure.

The ectodomains of Env trimers on immature virions are not expected to exhibit conformational changes that would prevent binding to CD4 and coreceptor. Indeed, immature virions can fuse with targets, although less efficiently than mature virions [62]. Consistent with Env trimers on immature virions binding to receptors on target cells and forming pre-hairpin intermediate structures, we occasionally found immature virions attached to target cells that were incubated with a fusion inhibitor. Immature virions comprised <5% of the datasets recorded. Attached immature virions exhibited comparable numbers and locations of spokes as we found for attached mature virions (Figure 4), consistent with a similar distribution of Envs on mature and immature virions. Although measurements of Env mobility within the membranes of mature and immature virions differ, the HIV-1 viral membrane on both mature and immature virions is a low mobility environment, likely due to high lipid order [61]. Our results suggest that the intrinsically low mobilities of HIV-1 Envs on both mature and immature virions limit the number of Env trimers that can participate in pre-hairpin intermediate structures with target cells.

The cytoplasmic tail of HIV-1 Env interacts directly with the viral matrix protein [63], and tail deletion has been suggested to increase the lateral mobility of Envs in the viral membrane [23, 24]. In order to determine the effects of cytoplasmic tail deletion on virion attachment to a target cell in the presence of a fusion inhibitor, we used ET to compare attachment sites for virions with wild-type and cytoplasmic tail-deleted Envs. We found a slight, but statistically significant, increase for the number of spokes in attachment sites formed between fusion inhibitor-treated target cells and the 6535-ΔCT versus wild-type pseudoviruses (3.9 +/- 1.2 for 6535-ΔCT to 2.4 +/- 0.5 for wild-type) (Figure 3d). These results are consistent with intrinsically low mobilities of HIV-1 Envs in the viral membrane that are increased only slightly for Envs that lack a cytoplasmic tail. Taken together, analyses of the numbers of apparent pre-hairpin intermediate structures trapped by fusion inhibitors on wild-type mature, wild-type immature, and cytoplasmic tail-deleted virions is consistent with the assumption that a minority of the relatively few HIV-1 Env trimers are involved in attachment via a pre-hairpin intermediate to target cells.

We also used the tomographic data to measure the dimensions at the virion-target cell attachment sites, finding that the approximate dimensions of the majority of the spokes (100-150 Å in length x 20-35 Å in width) are consistent with a pre-hairpin intermediate structure formed by gp41 alone after gp120 has dissociated (step iv, Figure 1a), which we measured to be ~100 Å x ~22 Å (Figure 1b). By contrast, a hypothetical pre-hairpin intermediate containing gp120 (step iii, Figure 1a) would be wider, with dimensions ~100 Å by ~120 Å (Figure 1b). This wider structure is roughly consistent with the ~100 Å x ~100 Å dimensions of each of the 5-7 rods of density that comprised the “entry claw” in a previous ET study [25]. We suggest that the entry claw rods visualized after incubating virus-target cell samples at 4°C and then warming to 37°C represent a structure formed after the HIV-1 Env trimer has bound to CD4 (and possibly also to a coreceptor) but prior to gp120 dissociation. Given that entry claws were not visualized in the temperature jump protocol when the viruses and target cells were incubated in the presence of a fusion inhibitor [25], the rod structures are unlikely to represent a pre-hairpin intermediate. We therefore suggest that the rods likely correspond to a structure resembling step ii in Figure 1a.

*A* potential explanation for why we observe 2-3 densities at virion-target cell attachment sites versus the 5-7 reported previously [25] could relate to the observation that the attachment surfaces in our imaging experiments were generally flat, as opposed to including a concave surface on the target cell that followed the convex surface of the virion [25]. Thus if HIV-1 virions contain a random assortment of a small number of Env trimers that diffuse slowly or not at all in the membrane, then only a small number of Envs would be available on the small contact surface formed by a roughly spherical virion and a flat cell membrane, whereas more Envs would be available for contacts with a larger contact surface formed by a virion interacting with a concave cell membrane (Figure 1c). Indeed, a recent ET study of murine leukemia virus (MLV) attached to target cells revealed a concave surface on the target cell with ~28 spokes per attached virion [64], a number that would be predicted to be higher than observed for HIV-1 even if HIV-1 were interacting with a concave portion of its target cell because MLV includes more spikes (at least 100 per virion) [65] compared with HIV-1 (7-14 per virion) [52–56].

In summary, we have used fusion inhibitors and ET of optimally preserved samples to visualize HIV-1 virions caught in the act of fusion with target cell membranes. These experiments revealed details of attachment sites with spokes representing pre-hairpin intermediate structures of HIV-1 Env. The spokes likely correspond to extended Env trimers after gp120 dissociation and prior to collapse into the post-fusion six-helical bundle structure (step iv in Figure 1a). Our observation of relatively few (2-3) spokes per attached virion implies that HIV-1 Env-mediated membrane fusion may require only a fraction of the ~14 Envs on each virion [52–56] and provides further details of an intermediate step on the pathway to viral entry.

## Methods

### Preparation of fusion inhibitors

A gene encoding T1249-Fc was constructed to encode the 39-residue T1249 peptide sequence [12] fused at its C-terminus to a (Gly4Ser)7 linker followed by human IgG1 Fc. T1249-Fc were expressed by transient transfection in 293-6E (CNRC) or Expi293 (ThermoFisher Scientific) cells and purified from transfected cell supernatants using a HiTrap MabSelect SuRe column (GE Healthcare) followed by size exclusion chromatography (SEC) using a Superdex 200 column (GE Healthcare), and its concentration was determined by A280 nm measurements using an extinction coefficient of 129,550 M^-1^ cm^-1^. D5 IgG was obtained from the NIH AIDS Reagents program at a stock concentration of 8.5 mg/mL. CPT31 was synthesized as described with its concentration measured by A280 (extinction coefficient of 37,980 M^-1^ cm^-1^) [13]. Z004 IgG, an anti-Zika virus antibody used as a control for non-specific entry inhibition, was expressed and purified as described [51].

### In vitro neutralization assays

SC4226618, 6535, and 6535-ΔCT (Env gene truncated after stop codon corresponding to gp41 residue Phe752) pseudoviruses were produced by cotransfection of HEK 293T cells with an Env expression plasmid and a replication-defective backbone plasmid [50]. In vitro neutralization assays were performed by measuring the reduction of HIV-1 Tat-induced luciferase reporter gene expression in the presence of a single round of pseudovirus infection in TZM-bl cells [50]. Inhibitors were evaluated using a 4-fold inhibitor dilution series (each concentration run in duplicate). Nonlinear regression analysis was used to derive IC_50_ and IC90 values, the concentrations at which half-maximal and 90% inhibition, respectively, were observed.

### Incubations for fusion inhibitor ET experiments

TZM-bl cells were plated and cultured as described [50] on carbon-coated, glow-discharged synthetic sapphire disks (3 mm diameter, 0.05 mm thickness; Technotrade International). Briefly, ~50,000 to 70,000 cells in 1 mL were seeded in each well and then replaced after a day with fresh media containing DEAE-Dextran (12 μg/mL). T1249-Fc, CPT31, or D5 IgG fusion inhibitor, each at a concentration of 130 ug/mL, was combined with pseudovirus (either SC4226618, 6535, or 6535-ΔCT, each at ~5000 TCID_50_/mL), and then immediately added to the cells and incubated for 2, 4, or 48 hours at 37°C. For control experiments, incubations were done with either no fusion inhibitor or with or with 130 μg/mL control IgG (Z004). In another control with SC4226618 pseudovirus, we added 0.01 μg/mL T1249-Fc fusion inhibitor (a concentration corresponding to 100-fold lower than the IC_50_ value for T1249-Fc against SC4226618). For all incubations, supernatant was then removed from each well, and cells were lightly fixed with 3% glutaraldehyde, 1% paraformaldehyde, 5% sucrose in 0.1 M sodium cacodylate trihydrate to render the samples safe for use outside of BSL-2 containment.

### EM Preparation and Electron Tomography

Sapphire disks were rinsed briefly with 1-Hexadecene (Sigma), placed individually into brass planchettes (Type A/B; Ted Pella, Inc.), and rapidly frozen with a HPM-010 High Pressure Freezing machine (BalTec/ABRA). Disks were transferred under liquid nitrogen to cryo-vials (Nunc) containing 2.5% OsO4, 0.05% uranyl acetate in acetone and then placed in a AFS-2 freeze substitution machine (Leica Microsystems, Vienna). Samples were freeze substituted at −90°C for 72 h, warmed to −20°C over 12 h, held at that temperature for 24 h, and then warmed to room temperature and infiltrated with Epon-Araldite resin (Electron Microscopy Sciences, Port Washington, PA). Sapphire disks were flat-embedded on teflon-coated glass slides with Secure-Seal adhesive spacers (Sigma) and Thermanox plastic coverslips (Electron Microscopy Sciences). Resin was polymerized at 60°C for 24 h.

Once embedded, the sapphire disks were removed, leaving the cells as a monolayer within the resin wafer. The cells were observed with an inverted phase-contrast microscope to ascertain preservation quality and to select regions of interest (i.e., regions with >10 cells in close proximity). These regions were extracted from the resin wafer with a microsurgical scalpel and glued to plastic sectioning stubs. Semi-thick (300-400 nm) serial sections were cut with a UC-6 ultramicrotome (Leica Microsystems, Vienna) using a diamond knife (Diatome, Ltd., Switzerland). Sections were collected onto formvar-coated copper-rhodium 1 mm slot EM grids (Electron Microscopy Sciences) and stained with uranyl acetate and lead citrate. Colloidal gold particles (10 nm) were placed on both surfaces of the grid to serve as fiducial markers for subsequent tomographic image alignment.

Grids were placed in a dual-axis tomography holder (Model 2040; E.A. Fischione Instruments, Export, PA) and imaged with a Tecnai TF-30ST transmission electron microscope (Thermo-Fisher Scientific) operating at 300 keV. Images were recorded with a XP1000 CCD camera (Gatan, Inc.). For dual-axis tomography, grids were tilted +/- 64° and images taken at 1° intervals. The grid was then rotated 90° and a similar tilt-series was taken about the orthogonal axis. Tilt-series datasets were acquired automatically using the SerialEM software package [66]. Tomograms were calculated, analyzed and modeled (including isosurface renderings) using the IMOD software package [67–69] on Mac Pro and iMac Pro computers (Apple, Inc.).

### Identification and imaging of HIV-1 virions

Prior to collecting tomographic data, HIV-1 virions were identified as follows: Thick sections were observed in the electron microscope and peripheries of cells were surveyed at medium magnification (3900x – 6500x). Objects that appeared to be spherical, were estimated to have a diameter of ~100 nm, and were proximal to a cell surface were examined at higher magnification (12,000x – 15,000x). These objects were observed at 0° tilt (perpendicular to the beam) and at +35° and −35° tilts to confirm that they were indeed spherical. Nonspherical objects, such as thin cellular projections or microspikes, would appear oblong or tubular at one or both high-tilt views. Objects that remained spherical were further evaluated by observing through a defocus series to detect core structures that would be indicative of a HIV-1 virion (i.e., a bullet-shaped mature core or a C-shaped immature core [40, 42]). Detection of core structures allowed the object to be classified as a HIV-1 particle, and it was subsequently imaged for dual-axis tomography. In most sample preparations, attached virions were found at an incidence of ~1 per every 5 cells in a given 400 nm section. On rare occasions, several (2-3) virions were found attached to a single cell, and each virion was recorded as a separate dataset. Spoke counts were determined from 3-D tomographic reconstructions (Figure 3d; Extended Data Table 1). Significance evaluations of spoke count differences between 6535-ΔCT and wild-type pseudoviruses was performed using a two-sample *t*-test assuming unequal variances.

## Acknowledgements

The authors thank Annie Lynch for assistance with initial experiments, Erica Lee and Jennifer Keeffe for preparing T1249-Fc and Z004 IgG, Sarah Apple and Nicholas Francis for preparing CPT31, Magnus Hoffmann for suggestions, and members of the Bjorkman and Kay laboratories for helpful discussions and critical review of the manuscript. This work was supported by the National Institute of General Medical Sciences (2 P50 AI150464) (to MSK and PJB). We thank the Caltech Kavli Nanoscience Institute for maintenance of the TF-30 electron microscope.

---

**Supplementary Figure 1.**
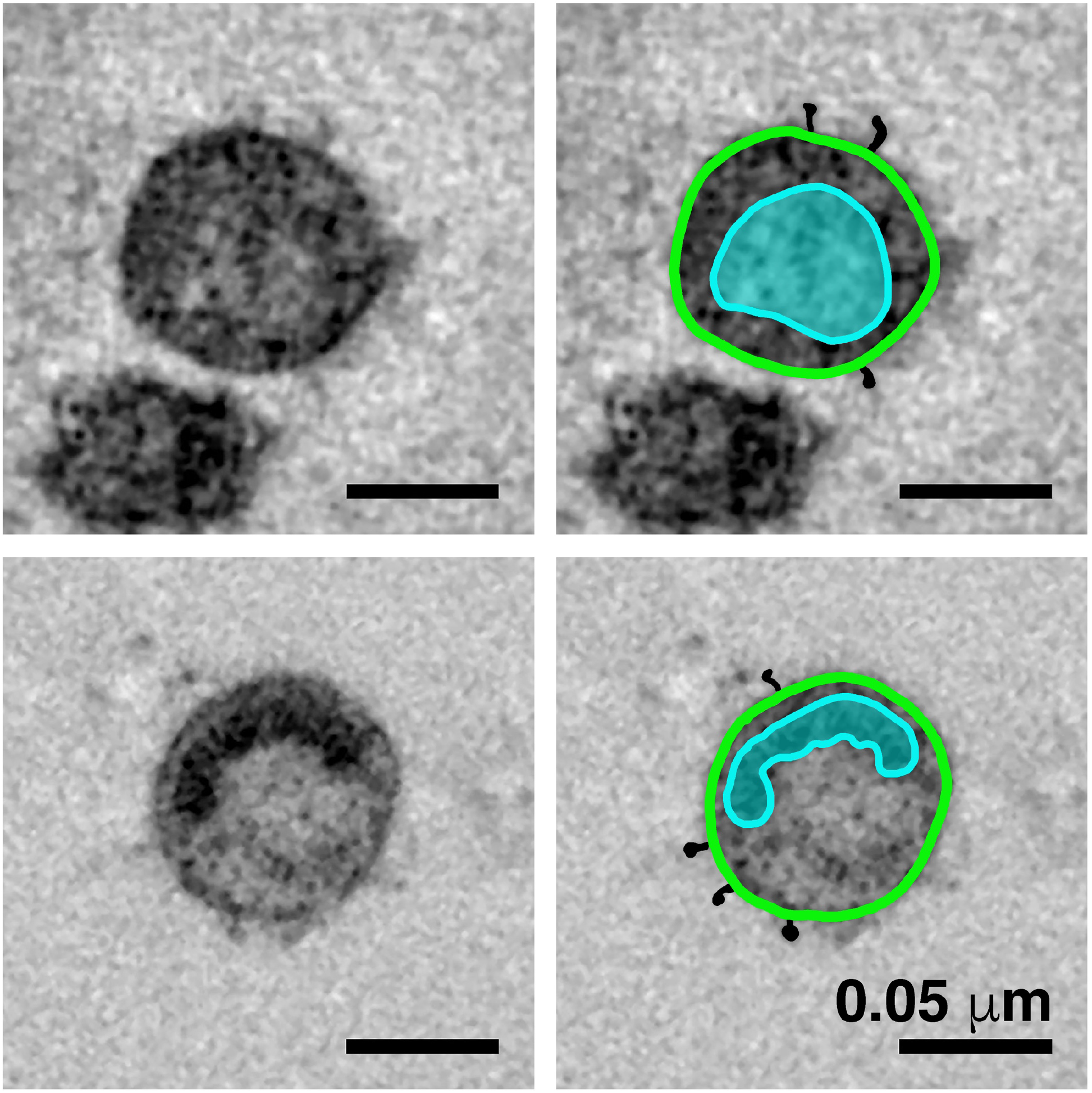
Identification of free pseudovirions in tomographic reconstructions. Virions were identified by visualizing a cone-shaped core structure in mature virions (upper left) and a C-shaped core structure in immature virions (lower left) and Env spikes on virion surfaces. Modeled contours highlight the core (blue) and Envs (black) in the mature virion (upper right) and in the immature virion (lower right).

**Supplementary Figure 2.**
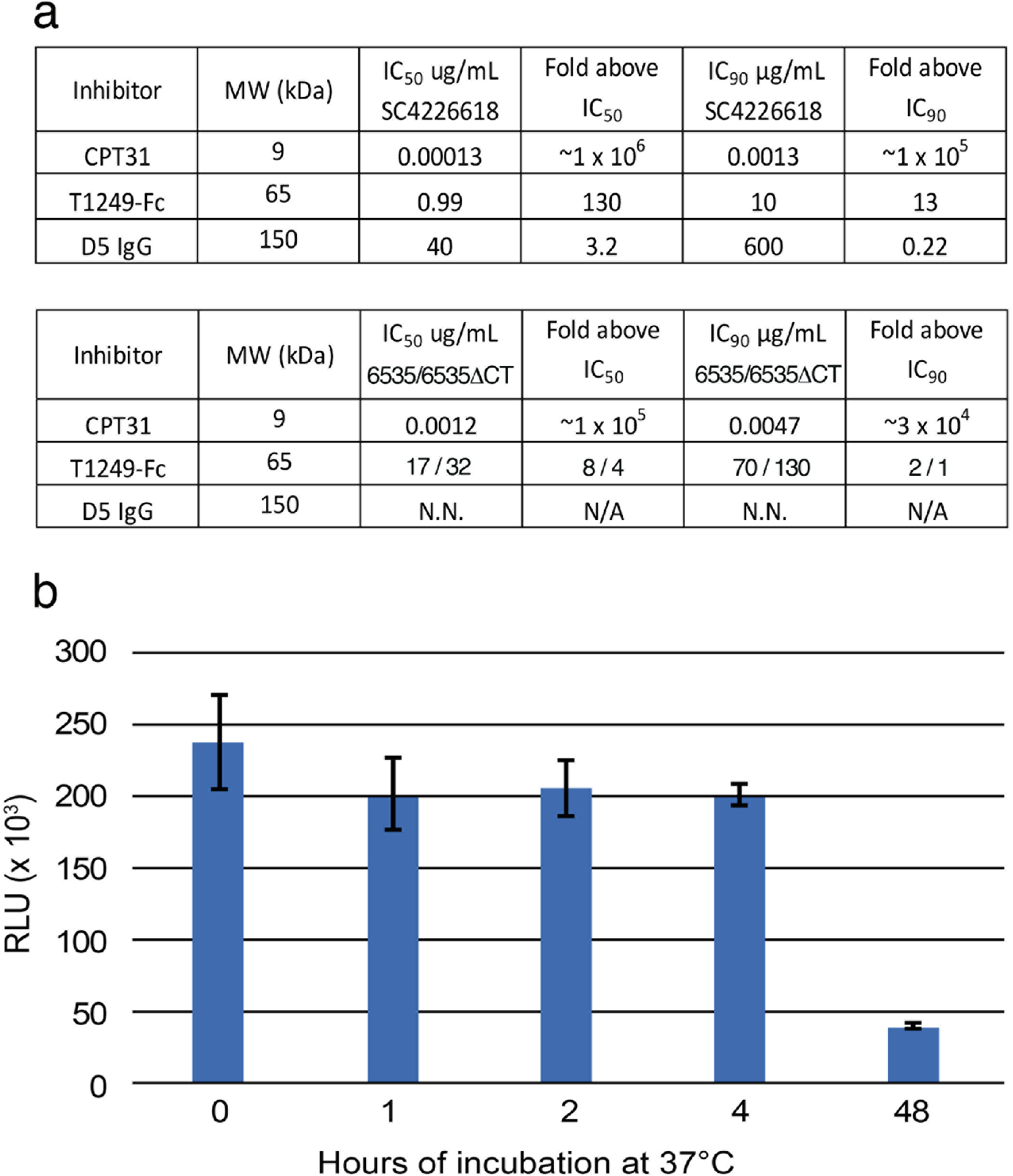
Characterization of fusion inhibitors and viral infectivity. **a**, Entry inhibitors are listed with their molecular weights, IC_50_ and IC90 values for neutralization potencies against SC4226618 and 6535 (and for T1249-Fc, also against 6535-ΔCT), and the fold above these values that they were used for fusion inhibitor imaging experiments in which the inhibitors were incubated at 130 μg/mL. N.N. = non-neutralizing. N/A = not applicable. **b**, Infectivity of SC4226618 pseudovirus after incubation with TZM-bl target cells for the indicated times at 37°C. Luciferase activity (measured in relative luminescence units, RLUs) of supernatants transferred to fresh TZM-bl cells after the following incubation times is presented as the mean and standard deviation for eight replicate measurements. ~20% of the input pseudovirus remained infectious after 48 hours at 37°C. To address whether infectious pseudovirus was still present after a 48-hour incubation at 37°C with T1249-Fc inhibitor and TZM-bl cells, we calculated how many molecules of T1249-Fc were added (10^15^ molecules) in the inhibition experiments, how many TZM-bl cells were cultured (~50,000 cells in each well, which contained 1 or 2 sapphire disks (3 mm each)), and how many SC4226618 pseudoviruses were added at the start of the incubation (2500 TCID_50_ = ~10^7^ particles). Thus there were ~2 x 10^6^ infectious viruses, ~10^15^ molecules of T1249-Fc, and ~50,000 TZM-bl cells that were available for formation of spoke structures that could be captured by HPF and visualized by ET after the 48-hour incubation at 37°C.

**Supplementary Figure 3.**
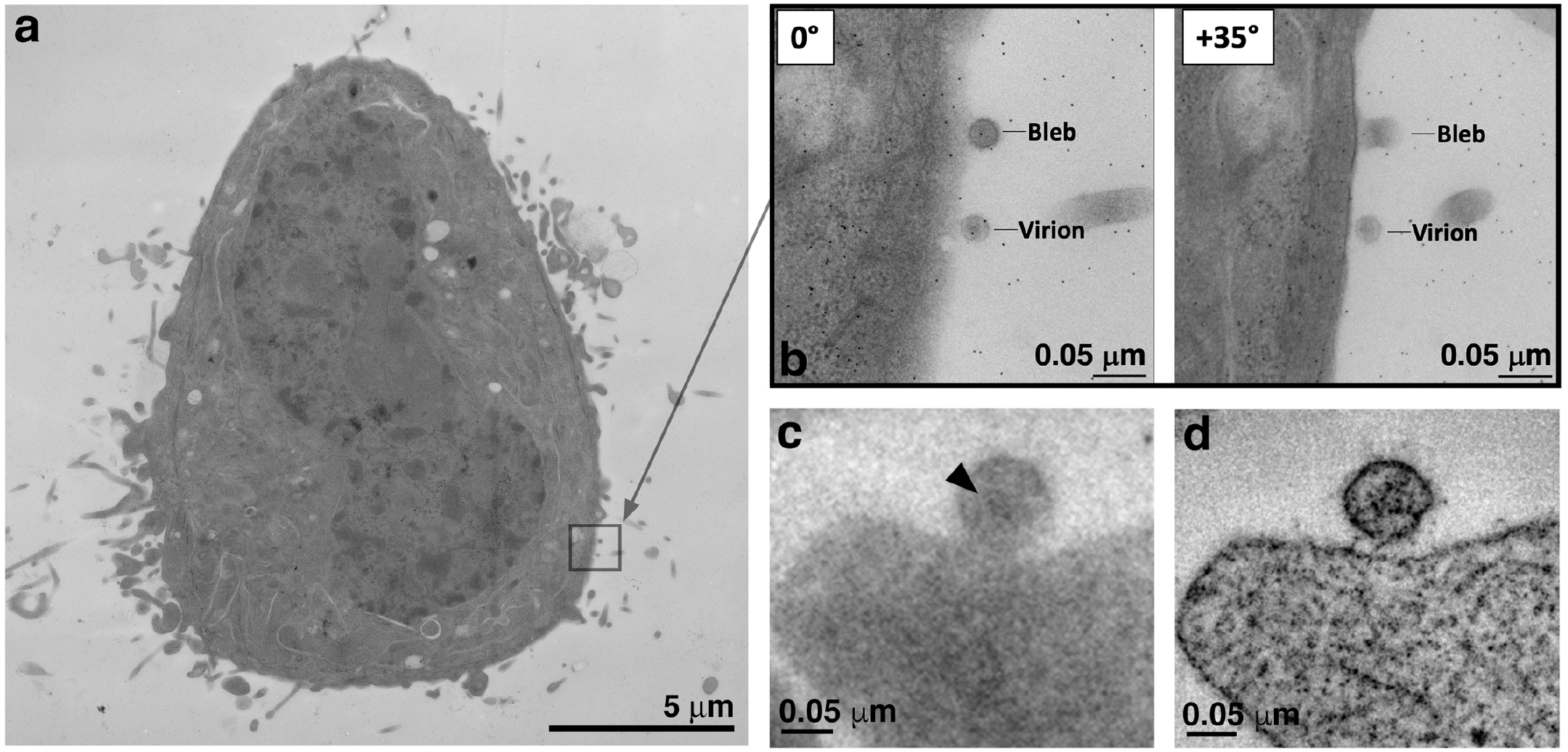
Locating and identifying attached virions prior to electron tomography. **a**, Overview of a TZM-bl cell in a 400 nm section. To find virions, the peripheries of cells are scanned at low magnification (2,600x – 5,600x) to identify structures that appear spherical, ~100 nm in diameter and proximal to the cell’s plasma membrane. Two candidate objects are indicated by the square. **b**, Tilted projection views of candidate objects. Once candidates are located, they are observed at high magnification (12,000x – 15,000x) first at 0° (view perpendicular to the electron beam) and then at +/-35°. Virions maintain a spherical appearance at both tilted views, while non-viral objects (e.g., blebs and other cellular extensions) appear tubular or oblong in the high tilted view. **c**, Projection image of a candidate virion at −5 μm defocus. Candidate objects are further examined at high magnification through a defocus series in order to distinguish core structures (a cone-shaped core in mature virions or a C-shaped core in immature virions). Objects with a distinguishable core (arrowhead) were classified as virions and imaged by dual-axis tomography. **d**, 5.5 nm slice from a tomographic reconstruction of the virion in panel c.

**Supplementary Figure 4.**
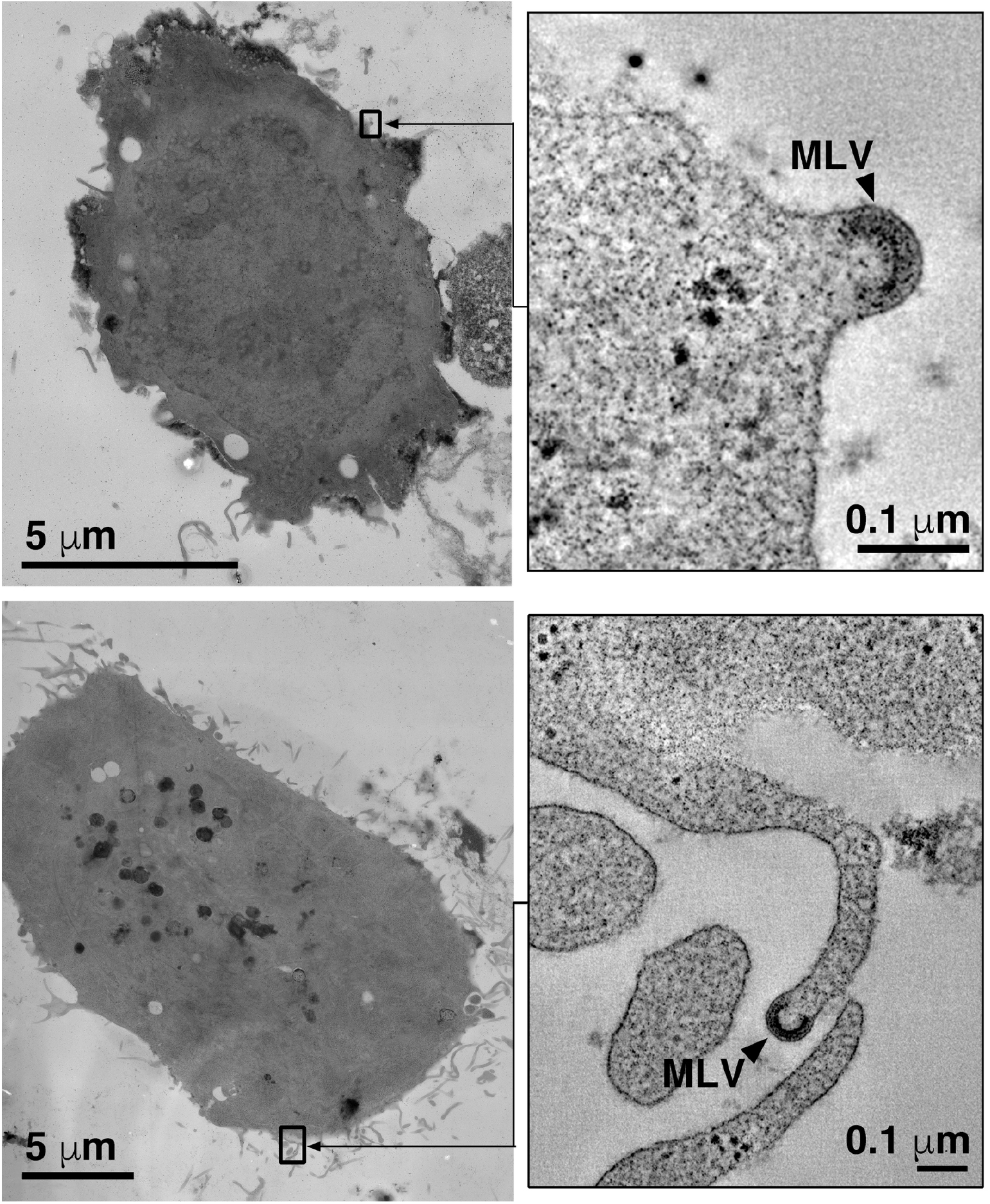
TZM-bl cells are contaminated with ecotropic murine leukemia virus. Two examples of nascent MLV particles emerging from the cell surface (top) and a cellular projection (bottom). Left panels show overviews of the cell with the locations of MLV budding events indicated by black rectangles. Right panels show tomographic details of the MLV budding profiles.

**Supplementary Figure 5.**
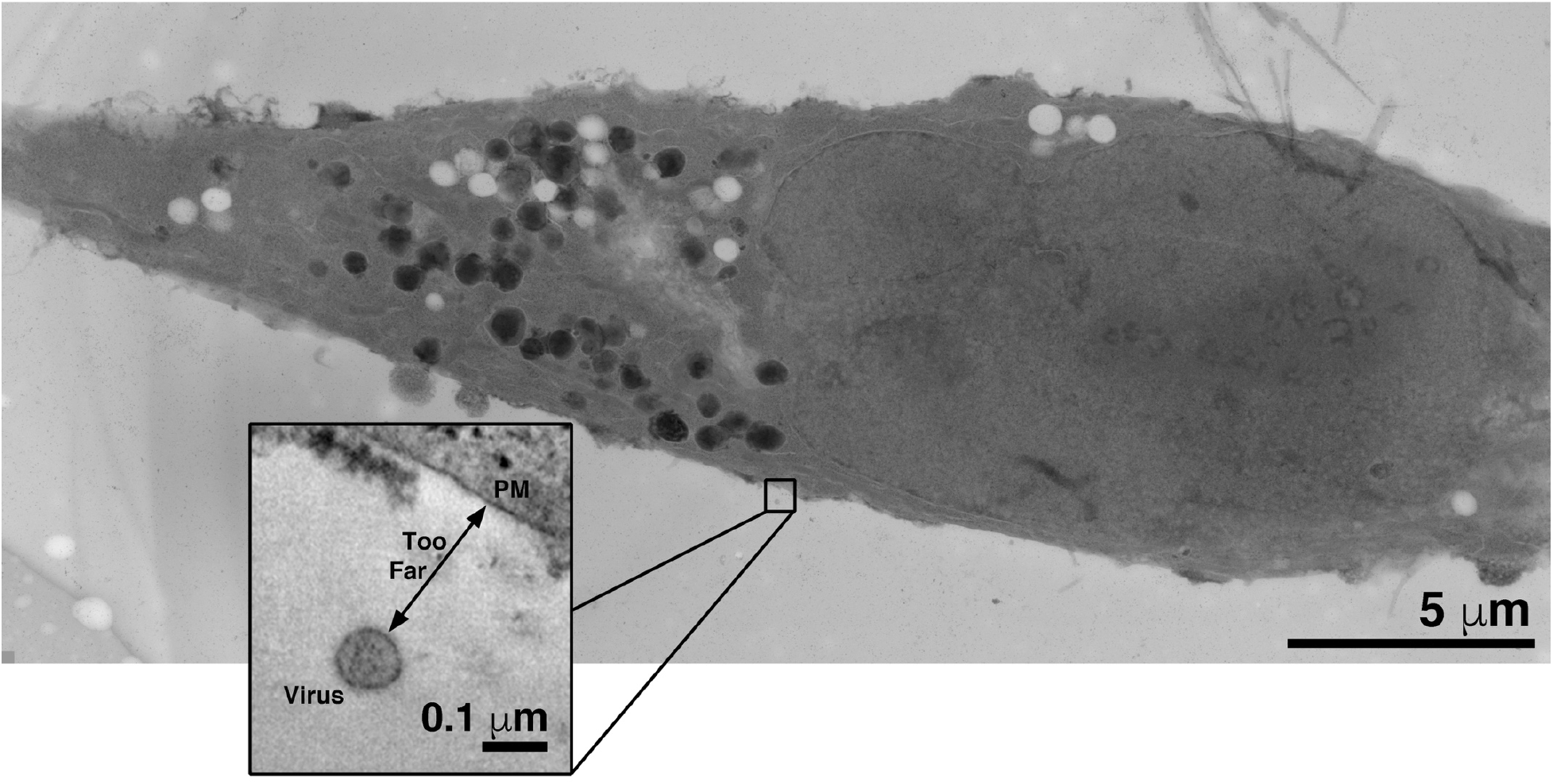
Control experiments showed no attached virions. No virions were found attached to cell surfaces when TZM-bl cells and pseudoviruses were incubated at 37°C with either no inhibitor, with an irrelevant Fc-containing protein (Z004, an anti-Zika virus IgG [51]), or with the T1249-Fc inhibitor at a concentration equivalent to 0.01x of its neutralization potency (i.e., its IC_50_ value). Very few free virions were present in the samples with fewer still in proximity to cells (<1 particle per ~10 cells per section). In the above typical example from a control experiment, a single free virion was found near a TZM-bl cell in a 400 nm section. Tomography (inset) revealed the virion to be too distant from the cell for attachment to occur and no structures resembling spokes on either the virion or the cell surface.

**Supplementary Figure 6.**
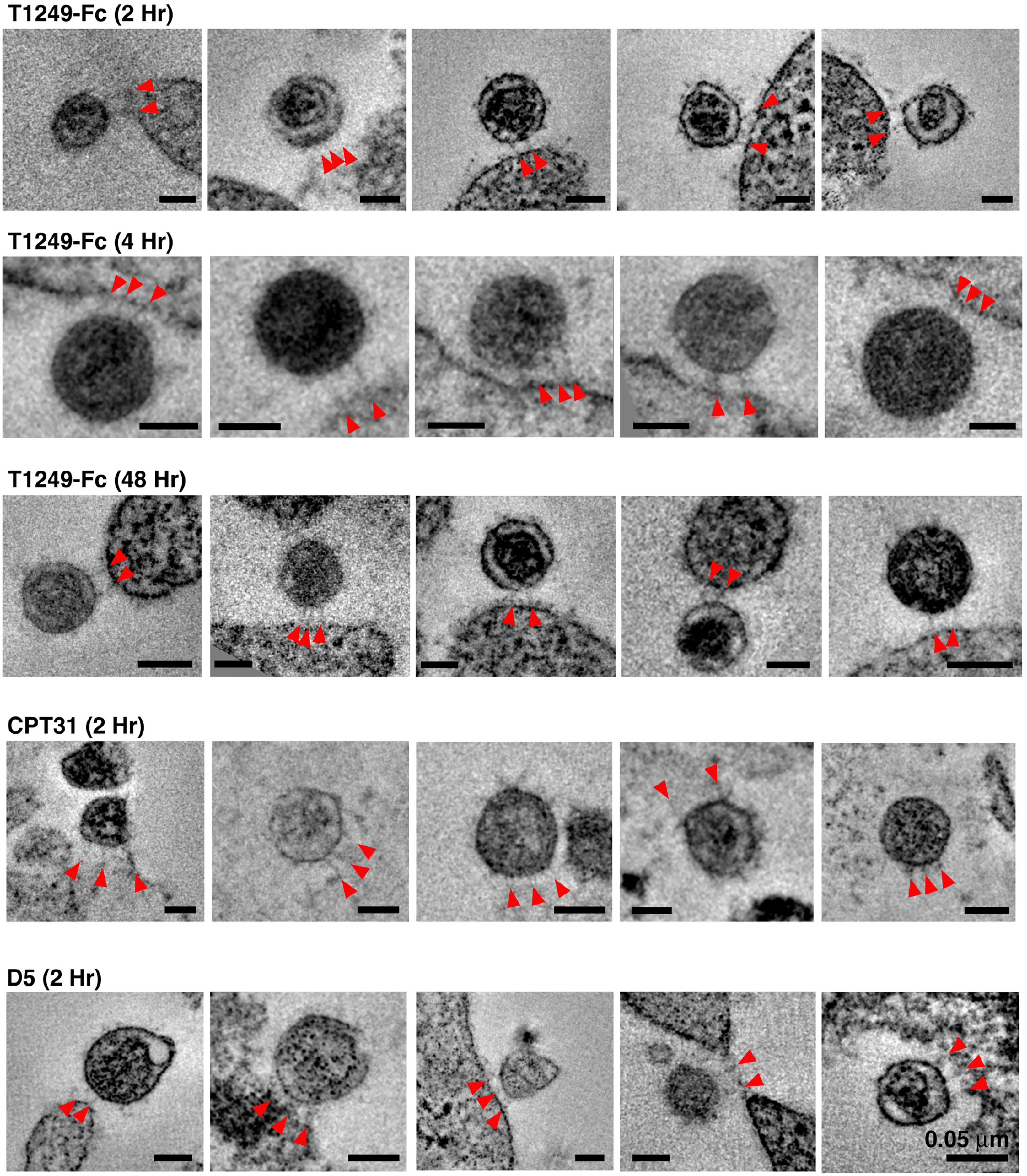
Gallery of attachment sites formed using different fusion inhibitors and different incubation times. In examples shown here, attachments consisted of either two or three spokes (red arrowheads) linking virions to cell surfaces. All scale bars = 0.05 μm.

## Supplementary Movie

Tomographic reconstruction of a mature HIV-1 pseudovirus (SC4226618) attached to a TZM-bl cell surface by two narrow spokes in an experiment in which the T1249-Fc inhibitor was incubated with cells and virus at 37°C for 2 hours. The movie presents the full volume of a 3-D reconstruction, advances at 1-pixel (0.5 nm) increments, and then pauses briefly to indicate the spokes (red arrowheads). The cone-shaped viral core is distinguished within the virion, as are Env trimers on the virion surface. Scale bar = 0.05 μm.

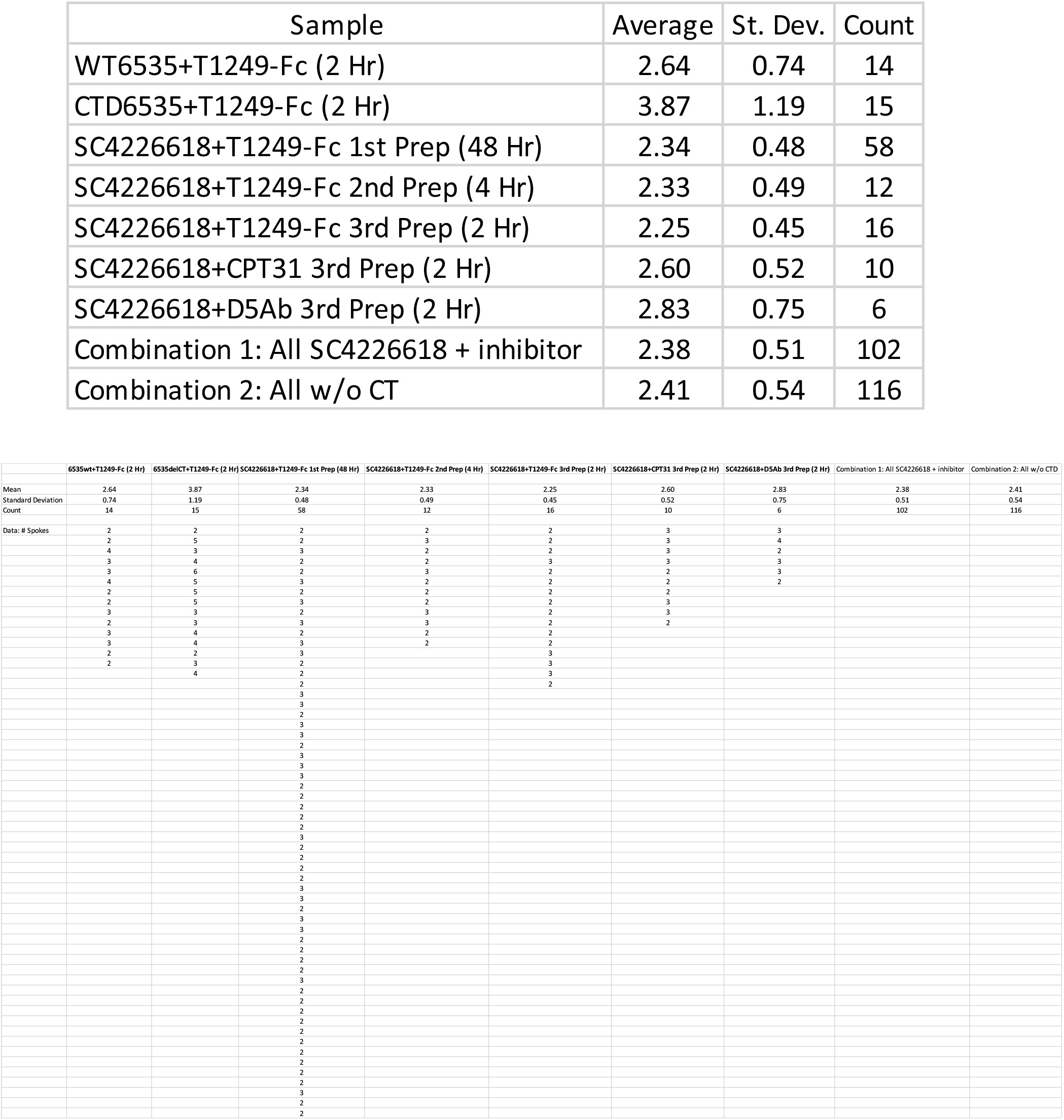
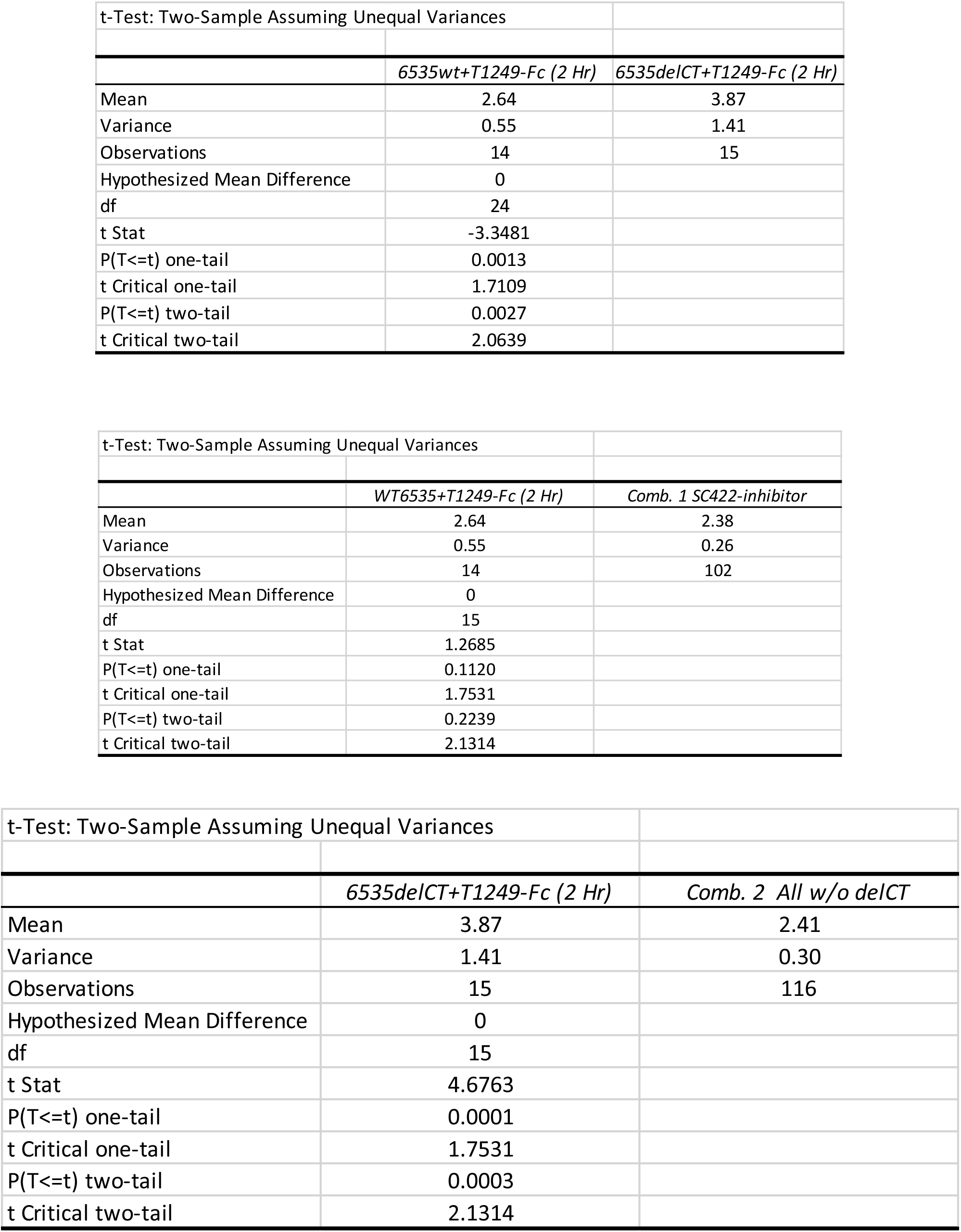

